# Pigment pattern morphospace of *Danio* fishes: evolutionary diversification and mutational effects

**DOI:** 10.1101/2021.05.10.443456

**Authors:** Braedan M. McCluskey, Yipeng Liang, Victor M. Lewis, Larissa B. Patterson, David M. Parichy

## Abstract

Molecular and cellular mechanisms underlying differences in adult form remain largely unknown. Adult pigment patterns of fishes in the genus *Danio*, which includes zebrafish, *D. rerio*, include horizontal stripes, vertical bars, spots and uniform patterns, and provide an outstanding opportunity to identify causes of species level variation in a neural crest derived trait. Yet understanding such variation requires quantitative approaches to assess phenotypes, and such methods have been mostly lacking for pigment patterns. We introduce metrics derived from information theory that describe patterns and pattern variation in *Danio* fishes. We find that such metrics used singly and in multivariate combinations are suitable for distinguishing general pattern types, and can reveal even subtle phenotypic differences attributable to mutations. Our study provides new tools for analyzing pigment pattern in *Danio* and potentially other groups, and sets the stage for future analyses of pattern morphospace and its mechanistic underpinnings.

**Summary statement:** We provide quantitative metrics for studying pigment patterns of zebrafish and other species. These metrics are applicable to changes between species as well as impacts of laboratory induced mutations

## Introduction

Elucidating the cellular and genetic bases for species differences in form remains a fundamentally important problem in biology. Considerable progress has been made in identifying allelic variants contributing to trait differences between populations of very closely related species through analyses of genetic crosses or naturally segregating variation. Yet it often remains unclear how differences in gene activity are translated through specific cellular behaviors of morphogenesis, differentiation, or both to generate particular morphological outcomes. In this regard, species closely related to developmental model organisms can be especially useful, as tools developed for mechanistic studies within species can often be adapted for testing hypotheses across species. With the advent of new methods of mutagenesis and transgenesis, many relatives of model organisms should become increasingly useful in comparative analyses.

One system proving its utility for elucidating mechanisms of morphological evolution is the diversity of adult pigment patterns in minnows of the genus *Danio* [1-3]. These patterns reflect the arrangements of pigment cells—black melanophores, yellow/orange xanthophores, and iridescent iridophores—that arise directly from neural crest cells, and indirectly from neural crest cells via latent precursors that differentiate during post-embryonic development [4-13]. The biomedical model organism zebrafish, *D. rerio*, develops horizontal dark stripes of melanophores and iridophores that alternate with light “interstripes” of xanthophores and iridophores. Yet even among the closest relatives of zebrafish [14] are taxa with horizontal stripes (*D. rerio, D. quagga, D. nigrofasciatus*), vertical bars (*D. aesculapii*), and spots (*D. kyathit, D. tinwini*). Elsewhere in *Danio*, sister species can exhibit a mostly uniform pattern (*D*. aff. *albolineatus*), wide stripes (*D. kerri*), bars (*D. erythromicron*) or “inverse” spots (*D. margaritatus*). Several additional patterns occur as well [15-21]. These pigment patterns influence shoaling behavior in the laboratory [22-25], and patterns of other fishes, and perhaps *Danio* species, function in mate recognition and mate choice, and can impact predation susceptibility [26-28].

*Danio* pigment patterns offer an outstanding opportunity to learn about the genetic and cellular mechanisms underlying trait evolution, given their largely two dimensional organization, tractable number of cell types, accessibility to imaging as phenotypes are developing, and approaches available for interrogating developmental genetic mechanisms and modeling pigment cell behaviors [3, 10, 29-38]. Given their developmental origins, these patterns may inform our understanding of species differences in other neural crest derived traits as well. To enhance the utility of this system, it will be important to develop methods for rigorously quantifying pattern variation across species and genotypes. Here, we introduce new metrics for assessing species patterns and phenotypes resulting from genetic perturbations. Our analyses lay the groundwork for future quantitative and mechanistic analyses of pattern diversification in this group.

## Results

Pigment patterns of small *Danio* species can mostly be classified as horizontally striped, spotted, or vertically barred. Understanding how these patterns arise and diversify is a necessary step in utilizing naturally occurring and laboratory-induced variants to understand pattern evolution more broadly. Based on ancestral state reconstruction, the most likely scenario to explain pattern diversity across these species is repeated evolution away from an ancestral pattern of either stripes or bars (**Fig. 1**; **Fig. S1A**). The predominant adult pattern amongst large *Danio species* (the outgroup of small *Danio* species), develops from juvenile stripes strikingly similar to zebrafish, which are subsequently modified into a chain-like patterns exemplified by *D. dangila* (**Fig. S1B**). Together, these findings suggest that a zebrafish-like striped pattern has been diversifying for millions of years to give rise to the variety of patterns present in the genus. A limited description of patterns into discrete classes, however, fails to encompass the breadth of pigment pattern diversity within this group and would be of limited use for quantifying intraspecific pattern variation.

**Fig. 1.**
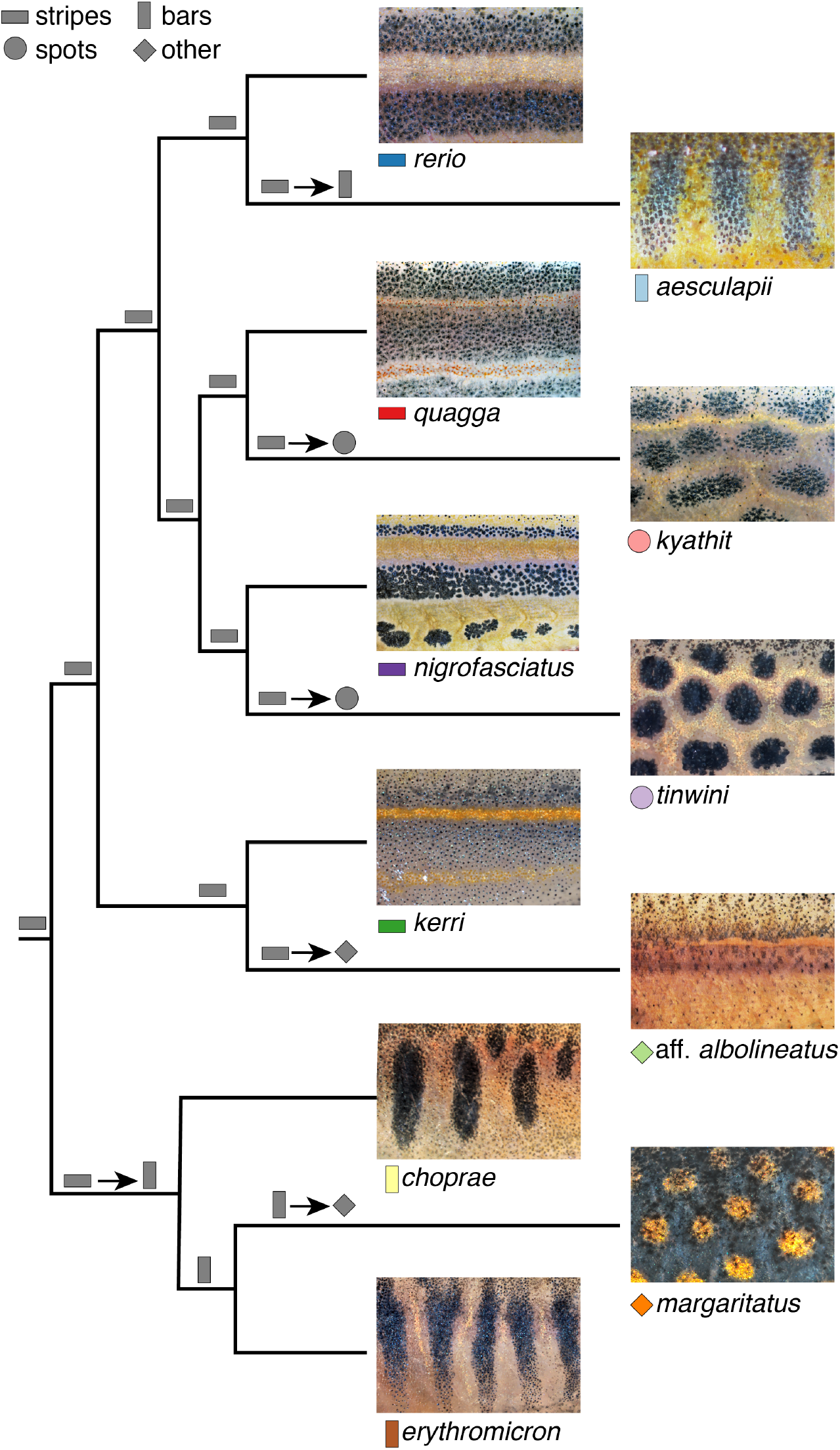
Diverse pigment patterns of *Danio* species. Shown are close-ups of pattern elements across eleven *Danio* species. Symbols next to species names denote overall pattern classes with colors and shapes correspond to those in Fig. 3. Gray shapes at nodes indicate the most likely pattern at that node based on ancestral state reconstruction (see Fig. S1) or preponderance of patten in outgroup species (root node). Shapes separated by arrows indicate transitions away from predicted ancestral patterns.

Quantitative descriptors of phenotypes and phenotypic variation are, therefore, critical for analyzing development, population variability and evolutionary diversification. Prior analyses of *Danio* pigmentation have relied principally on quantifications of cell numbers and positions relative to body axes, as well as nearest neighbor distances and binarized melanophore pattern elements [15, 31, 34, 38-40]. Because no single descriptor can fully characterize the complexity and variety of the patterns across *Danio* species and zebrafish mutants, we sought new metrics to facilitate mechanistic analyses and to enable a deeper understanding of pattern diversification, and particularly the morphospace within which development occurs [41]. We focused on gross features of patterns irrespective of color, and so captured images of several of the small *Danio* species that we converted to greyscale (**Fig. 2**). To make these images amenable to quantifications, we performed subsequent image processing steps (e.g., histogram standardization and gaussian blurring) using a pseudo-automated pipeline (**Fig. S2**) that accommodated minor technical differences in illumination intensity or quality, as well as biological variation associated with patterns themselves. To test the utility of our metrics for analyzing single locus mutant phenotypes, and to assess the roles of specific cell types in pattern formation, we additionally imaged fish with mutations that reduce or eliminate each major class of pigment cell owing to cell autonomous requirements of the affected gene: *colony stimulating factor 1 receptor a* (*csf1ra*) mutants, in which xanthophores are missing [39, 42, 43]; *KIT proto-oncogene, receptor tyrosine kinase a* (*kita*) mutants, in which melanophores are missing or reduced [40, 44, 45]; and *leukocyte receptor tyrosine kinase* (*ltk*) mutants, in which iridophores are missing [46, 47].

**Fig. 2.**
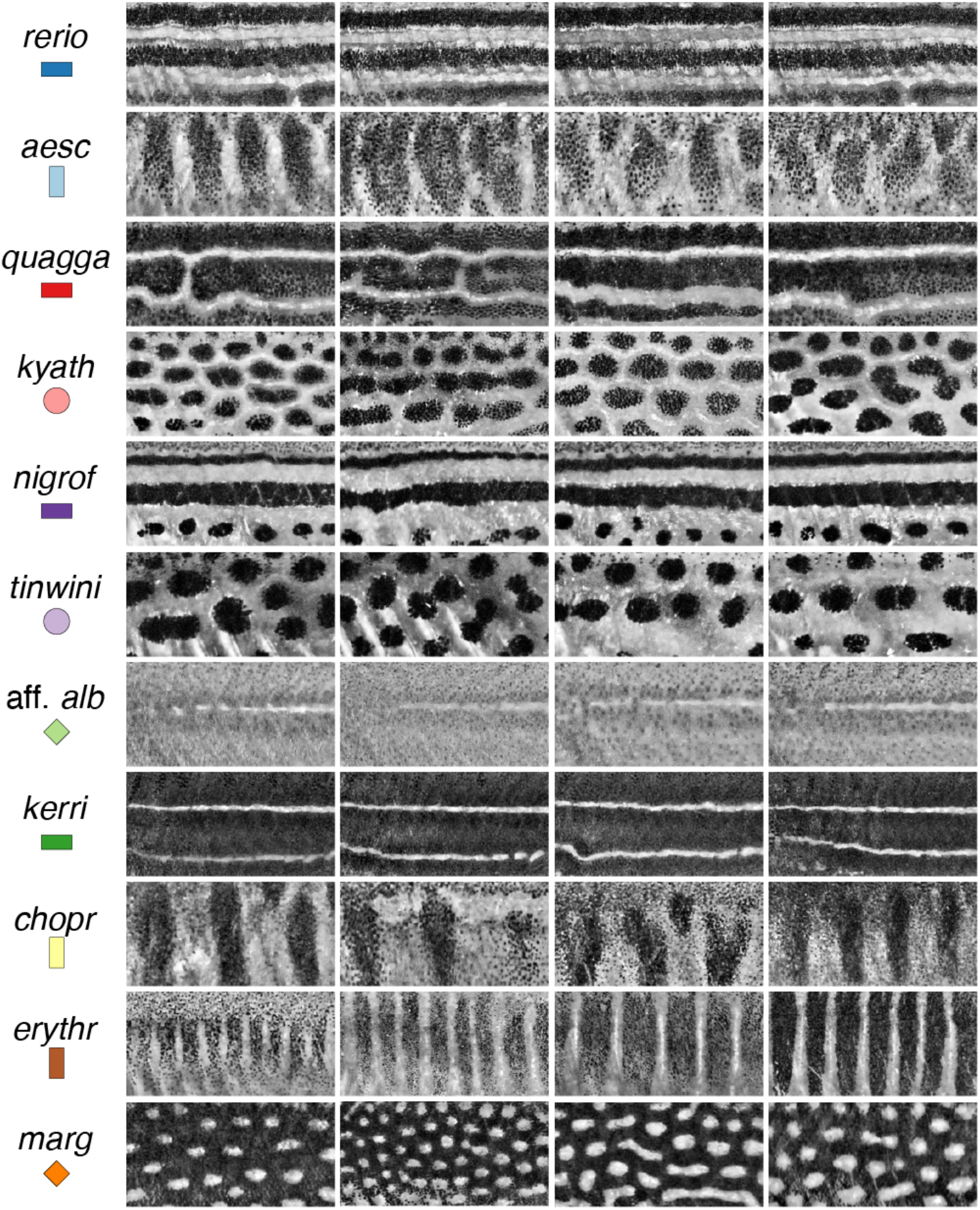
Species pattern variation. Panels show typical wild-type phenotypes within species and illustrate the regions of interest and greyscale conversions used for analyses. Left two columns show females, right two columns show males.

We first examined spatial entropy as a measure of pattern quality for wild-type fishes. Derived from information theory, entropy measures uncertainty [48], and, in this context, the average magnitude of shading differences along a spatial axis [24]. As entropy and similar metrics do not consider overall brightness, dark spots on a light background (as in *D. kyathit*) and light spots on a dark background (as in *D. margaritatus*), can have similar values despite having near opposite pixel intensities. **Figure 3A** shows mean entropies of individual fish along their dorsoventral (DV) axis (“stripe entropy,” *E*_*DV*_) and their anteroposterior (AP) axis (“bar entropy,” *E*_*AP*_). The plot defines a pattern morphospace with regions corresponding to stripes (*D. rerio, D. nigrofasciatus*), bars (*D. aesculapii, D. erythromicron, D. choprae*), and spots (*D. tinwini, D. kyathit, D. margaritatus*; **Fig. S3A,B**). Species with indistinct patterns or very narrow interstripes (*D*. aff. *albolineatus* and *D. kerri*) had low entropy values. Discriminant analysis using *E*_*DV*_ and *E*_*AP*_ scores (**Table S1**) classified 64% of individuals correctly to species, with misclassifications limited to species having similar patterns in bivariate space (e.g., *D. choprae, D. erythromicron*).

**Fig. 3.**
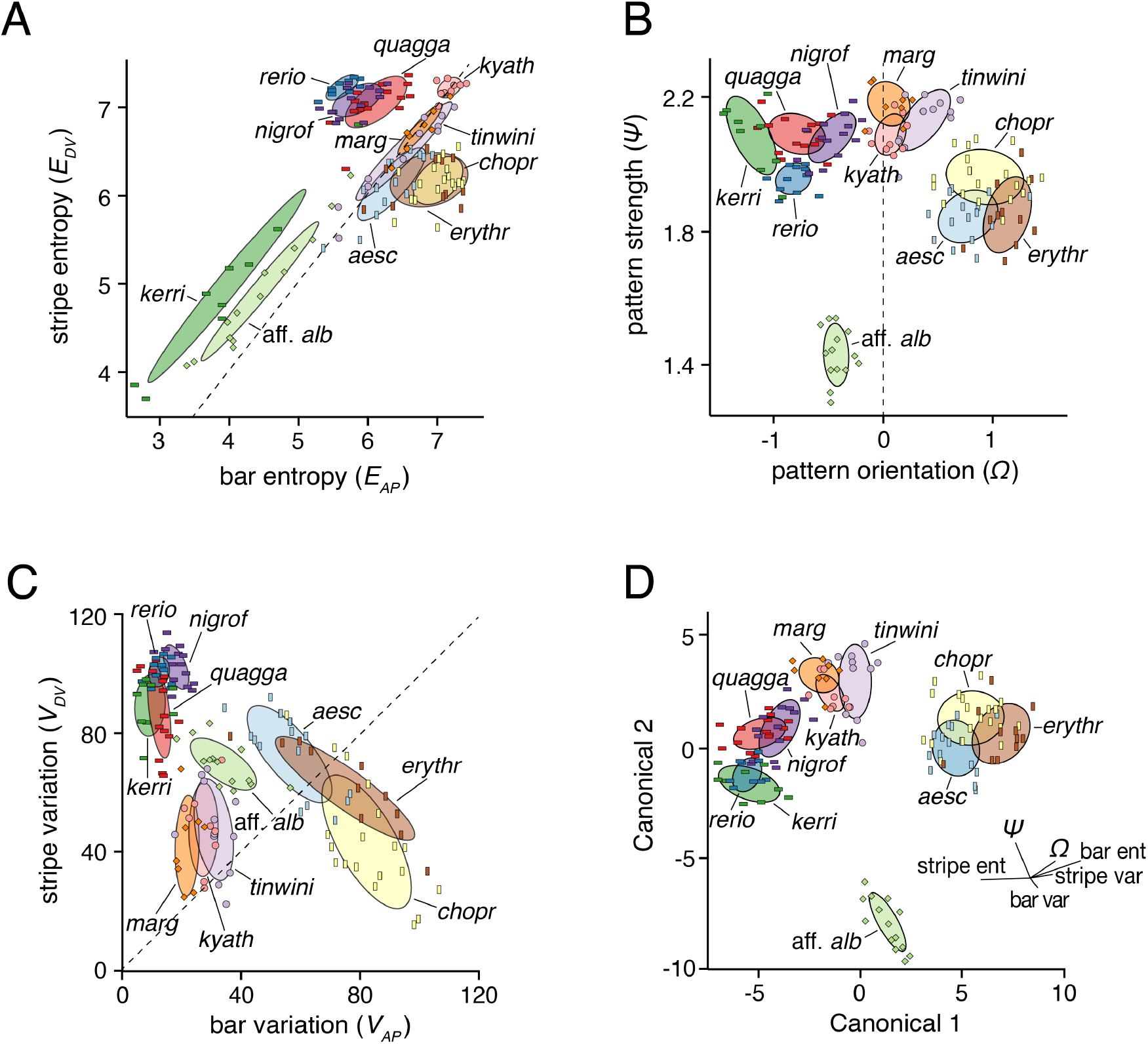
Metrics for describing pattern variation. (A) Entropies for patterns of individual wild-type fish measured along the DV axis (stripe entropy, *E*_*DV*_) and AP axis (bar entropy, *E*_*AP*_) with each point representing the mean entropies across transects within a single fish. Species symbols correspond to Figure 1. Dashed line denotes equal stripe and bar entropy. The plot reveals broad pattern types (e.g., stripes above dashed line, bars right of dashed line) and species-differences in pattern space. (B) Pattern strength (*ψ*) and orientation (*Ω*) also distinguished among pattern types (e.g., bars at right, stripes at left). Dashed line denotes patterns with similar horizontal and vertical contributions. (C) Species patterns are more dispersed in the morphospace defined by stripe variation (V_*DV*_) and bar variation (V_*AP*_) compared to the other morphospaces. Dashed line denotes equal stripe and bar pattern. (D) Species are best delimited in a higher dimensional space derived from all six pattern metrics. Shown are the species plotted against the first two canonicals resulting from a discriminant analysis using all six pattern metrics. Loadings of metrics along the first two canonicals are shown.

To identify additional pattern metrics, we converted two-dimensional spatial data to a frequency domain using two-dimensional fast Fourier transformation (FFT) [49-52]. We defined pattern orientation (*Ω*) to indicate the relative contributions of peaks in frequency between axes (DV:AP), distinguishing stripes (*Ω* < 0) and bars (*Ω* > 0), from spots or uniform patterns that lack orientation (*Ω* ≈ 0). To further capture variation in the distinctiveness of elements within patterns, we defined a metric of pattern strength (*ψ*) as the overall magnitude of peaks in frequency. **Figure 3B** plots *Ω* and *ψ*, illustrating pattern spaces corresponding to horizontal stripes (*D. quagga, D. rerio, D. nigrofasciatus*) and relatively weaker vertical bars (*D. erythromicron, D. aesculapii, D. choprae;* **Fig. S3C,D**). In the vicinity of *Ω* = 0, where patterns were relatively lacking in orientation, *ψ* distinguished strong patterns of highly contrasting elements (spots of *D. kyathit, D. margaritatus*) from weak patterns of a more diffuse nature (*D*. aff. *albolineatus*). Discriminant analysis using *Ω* and *ψ* classified 72% of individuals correctly to species, though patterns of *D. choprae* and *D. erythromicron* remained broadly overlapping in multivariate space.

For a third pair of metrics, we quantified the same images using a method designed around binarized patterns, segmented simply into melanized elements vs. non-melanized elements. This method has been useful for describing a transition between the striped pattern of *D. quagga* and the spotted pattern of *D. kyathit* [38]. Yet it remained unclear how effective the approach might be in describing patterns with less distinctive elements (e.g., *D*. aff. *albolineatus*) or cells that are more heterogeneously dispersed, as in *csf1ra, kita*, or several other mutants of *D. rerio* [39, 45, 53-57]. Following pattern segmentation, this method quantifies pattern variation along the dorsoventral (DV) axis (*V*_*DV*_, or “stripe variation”) and the anteroposterior (AP) axis (*V*_*AP*_, or “bar variation”). In this morphospace, striped species (*D. rerio, D. nigrofasciatus, D. quagga*) and barred species (*D. erythromicron, D. choprae*) fell at the high ends of their respective axes, while spotted species (*D. tinwini, D. kyathit, D. margaritatus*) clustered near the origin (**Fig. 3C**; **Figs. S3E,F**). Horizontally striped species each occupied relatively small areas of morphospace, whereas vertically barred species occupied relatively larger areas. This observation points to the inherently more stereotyped patterns of the former species, and the greater variability in pattern among individuals of the latter species (**Fig. 2**). Despite considerable overlap across species in this morphospace, discriminant analysis using *V*_*DV*_ and *V*_*AP*_ classified 67% of individuals to the correct species.

Assessing more than two metrics at a time allowed for an increased parsing of *Danio* morphospace and improved separation of species within it. Discriminant analyses using all six metrics simultaneously (*E*_*DV*_, *E*_*AP*_, *Ω, ψ, V*_*DV*_ and *V*_*AP*_) classified 96% of individuals correctly to species, demonstrating the benefit of incorporating distinct metrics to describe naturally occurring pattern variation. Plotting species according to the first two canonical variables derived from this analysis (**Table S1**) revealed areas of morphospace that clearly correspond to regions of striped, spotted, and barred patterns, with the uniformly patterned *D*. aff. *albolineatus* far removed from all other species (**Fig. 3D**). Additional canonical variables allowed for a finer discrimination of similar patterns across species (**Fig. 4**). Bivariate correlations between pattern metrics (**Table S2**) revealed information contents that are partly, though not entirely, overlapping. Overtly subtle but statistically significant differences in some metrics between sexes were also apparent (**Table S3**).

**Fig. 4.**
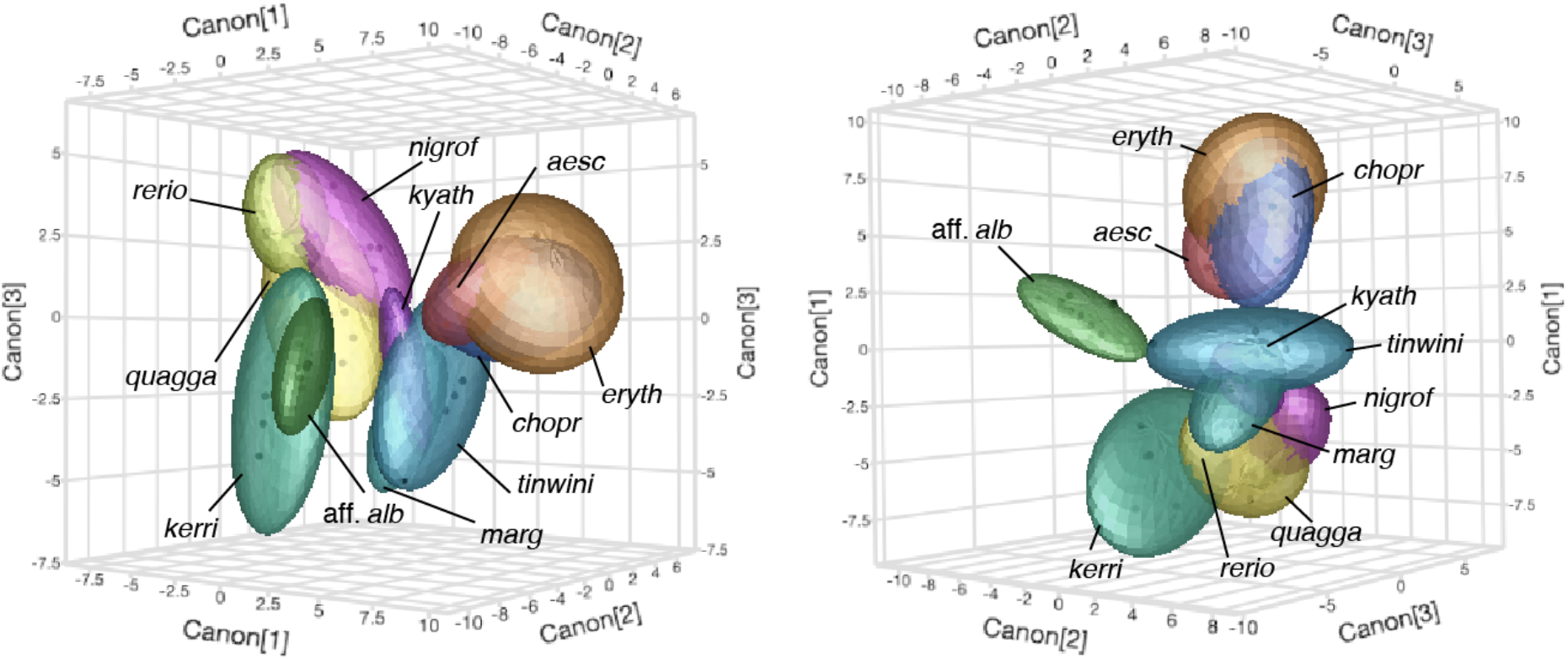
Multivariate pattern space separates pattern types and species. Shown are two views of pattern space represented in three-dimensions, as described by combinations of six pattern metrics (*E*_AP_, *E*_DV_, *Ω, ψ, V*_AP_, *V*_DV_). Bubbles show 95% confidence intervals. Discriminant analyses indicated significant predictive power for each metric (all *P*<0.0001) and classified 96% of individuals correctly to species.

To further assess the utility of these metrics for describing pattern differences resulting from induced or naturally segregating allelic variation, we examined *csf1ra, kita*, and *ltk* mutant phenotypes, deficient in xanthophores, melanophores and iridophores, respectively. Entropy, FFT, and binarized-variation metrics each revealed significant differences between wild-type and mutant patterns of each species (**Fig. S3**). The effects of some of these mutations varied between species and could best be summarized by comparing overt phenotypes (**Fig. 5A; Fig. S4**) to metrics describing pattern orientation and strength (**Fig. 5B–D**).

**Fig. 5.**
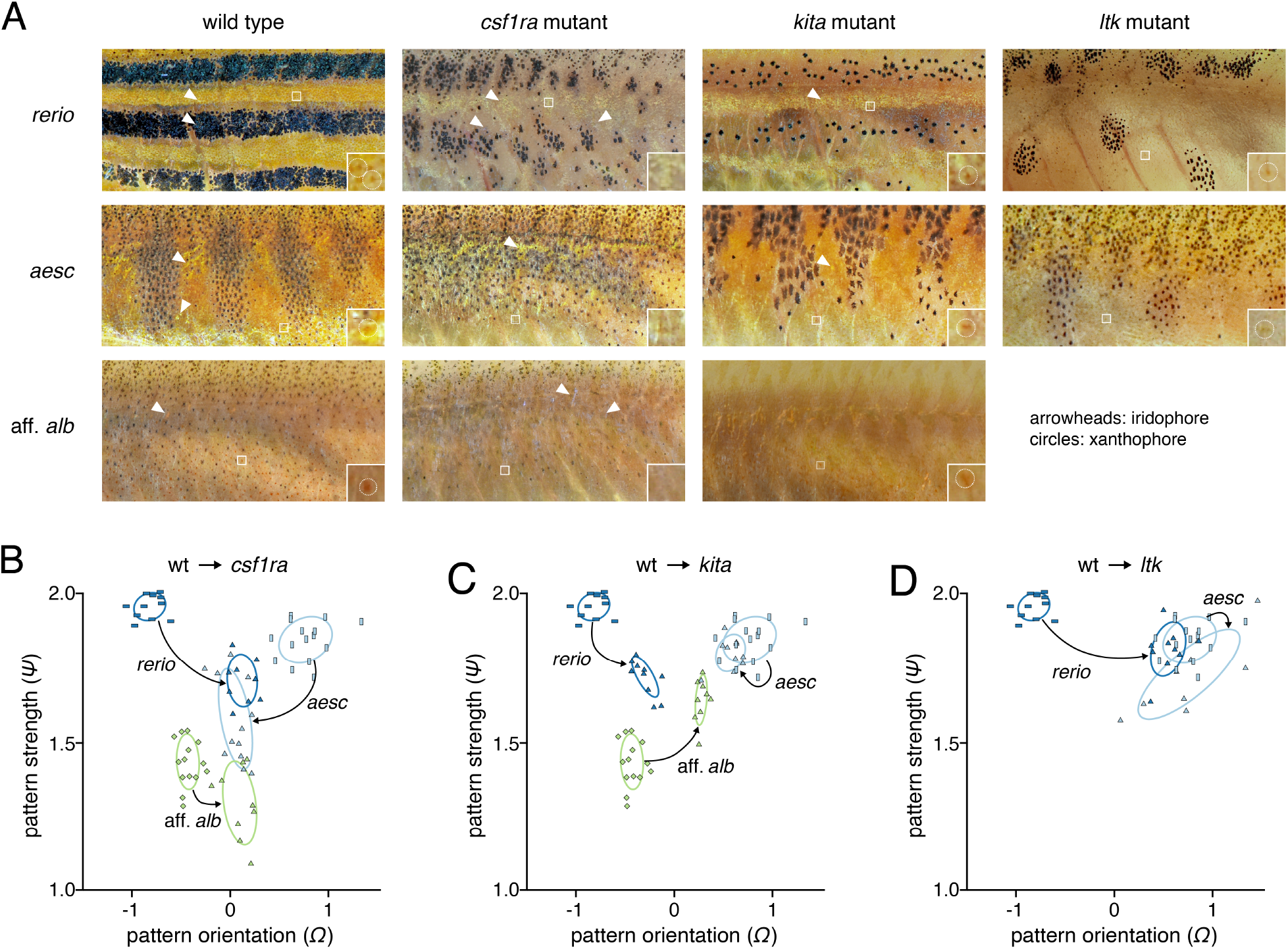
Effects of *kita, ltk*, and *csf1ra* mutation across species. (A) Pattern phenotypes of wild-type and mutant *D. rerio, D. aesculapii*, and *D*. aff. *albolineatus*. (B) Effects of *csf1ra* mutation on pattern strength and orientation in three *Danio* species. (C) Effects of *kita* mutation on pattern strength and orientation in three *Danio* species. (D) Effects of *ltk* mutation on pattern strength and orientation in two *Danio* species.

Removing xanthophores through *csf1ra* mutation reduced pattern strength and removed most directionality from the pattern (*Ω* ≈ 0) in all three species (**Fig. 5B**). Interestingly, *csf1ra* mutant *D. rerio* and *D. aesculapii* appeared to converge on the pattern of wild-type and *csf1ra* mutant *D*. aff. *albolineatus* in bivariate space, indicating a substantial dependence of periodic pattern on xanthophores in both striped and barred species (**Fig. 5B**), presumably owing in large part to interactions between these cells and pigment cells of the other classes [9, 30, 31, 39, 43, 47, 58, 59]. Notably these analyses detected even a subtle change in the pattern of D. albolineatus owing to the loss of xanthophores, such than an already nearly uniform pattern became even more so. This difference presumably reflected a more homogenous appearance of melanophores across the flank in the absence of xanthophores, even adjacent to the residual interstripe where these cells often have a darker, more spread appearance in the wild-type, particularly evident in greyscale images (e.g., **Fig. S4**).

In contrast to the shared loss of xanthophores and patterning consequences owing to *csf1ra* mutation, each species responded differently to *kita* perturbation (**Fig. 5C**). *kita* mutants of *D. rerio* retain some melanophores and develop a rudimentary stripe pattern [44], but had lower pattern strength and reduced directionality compared to their wild-type counterparts. *kita* mutants of *D. aesculapii* had fewer melanophores, but still formed rudimentary bars, which fell very close to the patterns of wild-type. By contrast, *kita* mutants of *D*. aff. *albolineatus* lacked virtually all melanophores (a phenotype more severe than observed previously for *kita* mutants of *D. albolineatus* [40]), leaving almost no pattern to quantify. This led to a position in morphospace lacking in orientation, yet aberrantly increased in strength presumably owing to the increased contrast of features beneath the skin without a covering of melanophores.

Losses of iridophores owing to mutations in *ltk* also had species-specific effects. In *D. rerio*, iridophores are essential for the normal establishment of pattern and normal numbers of melanophores, through their interactions with these cells and xanthophores; in the absence of iridophores interactions between residual melanophores and xanthophores generate a late-appearing, regulative pattern of several spots [31, 34, 47, 60, 61]. This alteration was reflected in a marked shift in morphospace (**Fig. 5D**). By contrast, the elimination of iridophores from *D. aesculapii* had only a limited effect on major pattern features, though mutant individuals tended to be more variable in appearance leading to them occupying a larger area of morphospace than wild-type. The persistence of pattern observed for iridophore-free *ltk* mutants was also consistent with the phenotype of *D. aesculapii* that lack iridophores due to mutation at a different locus (*mpv17*) [37].

Finally, because wild-type pattern elements, pattern-forming cell behaviors, rates and directions of growth, and effects of induced mutations can differ between anatomical regions [3, 13, 54, 62, 63], we asked whether metrics might be useful for describing pattern variation within individuals. Inspection of xanthophore-free *csf1ra* mutants of *D. rerio* suggested a difference between regions in the magnitude of effects on melanophore patterning, with a seemingly less severe defect in stripes anteriorly and more severe defect posteriorly (perhaps accounting in part for different interpretations of the importance of xanthophores in stripe formation [8, 39, 43, 64, 65]). Accordingly, we predicted the stripe entropies (*E*_*DV*_) should differ between anterior and posterior regions of the same individuals. Indeed, comparison of *E*_*DV*_ scores revealed a significant reduction in *csf1ra* mutants compared to wild-type, and this loss was more severe posteriorly than anteriorly (**Fig. 6**).

**Fig. 6.**
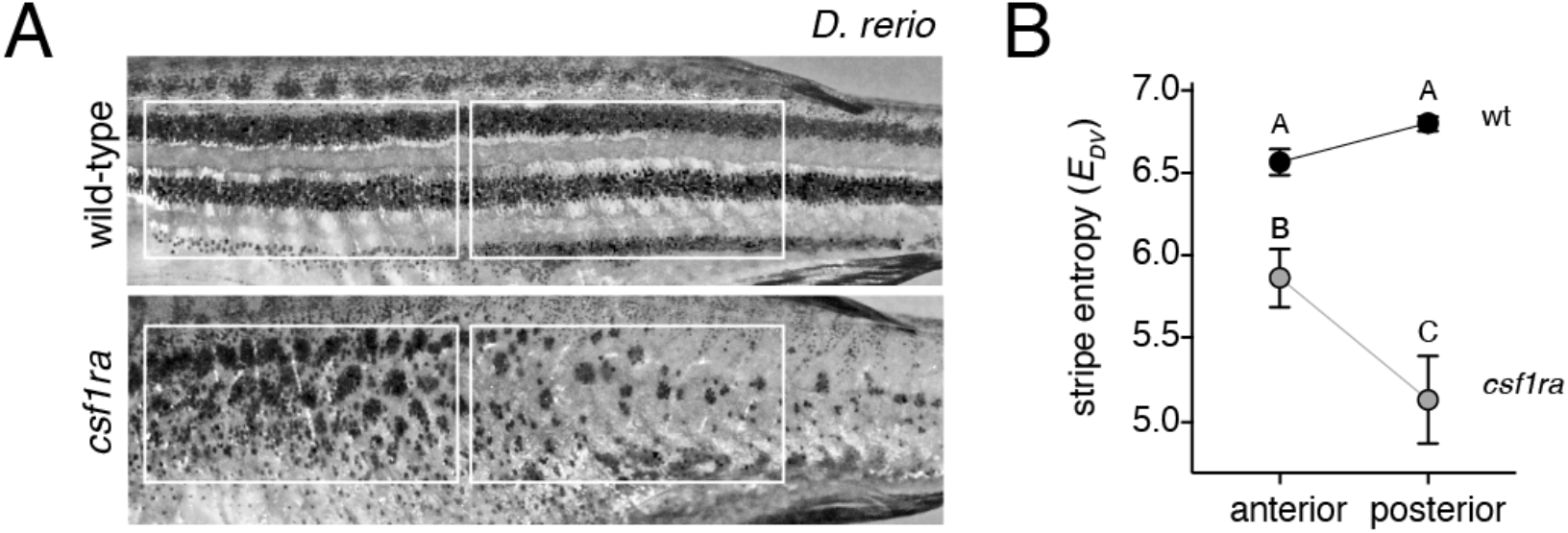
Entropy scores reveal anterior–posterior differences in *csf1ra* mutant phenotypes. (A) Representative wild-type and *csf1ra* mutant *D. rerio* illustrating regions of interest. (B) Stripe entropy scores (mean±SE) did not differ between anterior and posterior regions of wild-type, were reduced in *csf1ra* mutants overall, but especially in the posterior (genotype x region interaction, *F*_1,36_=16.6, *P=*0.0002; reciprocal values pf *E*_*DV*_ were analyzed to control for differences in residual variance among groups). Shared letters above symbols indicate means not significantly different from one another (*P*>0.05) in Turkey-Kramer post hoc comparisons.

## Discussion

This study provides metrics for understanding pattern development and evolution. We found that spatial entropy, variables derived from FFT analyses, and measurements of variation in melanophore element patterns were effective in describing patterns and discriminating among major pattern types (i.e., stripes, bars, spots, and indistinct patterns). Indeed, these metrics revealed even subtle differences between wild-type and *csf1ra* mutants in *D*. aff. *albolineatus*. Despite the utility of these metrics, several caveats apply. Correlations among some variables (**Table S2**) indicate information contents that are partly, though not entirely, overlapping. Moreover, our analyses suggest the need for additional independent metrics: even with six metrics, patterns of barred species (*D. aesculapii, D. choprae* and *D. erythromicron)* were broadly overlapping in multivariate space (**Fig. 4**), despite the fish being rather obviously different from one another (**Fig. 2**). Although discrepancies between quantitative outcomes and intuition could be biologically meaningful [24], these differences seem likely to reflect methodological vagaries, including regions used for comparison, and whether patterns of different species have different requirements for optimal illumination. Our metrics also do not inherently distinguish between pattern “features” and “backgrounds,” as dark spots on a light background (*D. kyathit, D. tinwini*) clustered with lights spots on a dark background (*D. margaritatus*). Likewise, the use of grayscale images elides differences in color that are themselves interesting, and particularly important to fish, even in spectral ranges not visible to the human eye [27, 66]. Capturing these and other phenotypic attributes will require the development and incorporation of additional metrics. Nevertheless, the approaches we present offer a quantitative framework for comparing relative impacts of mutations on diverse patterns, will facilitate the determination of whether some patterns are more or less likely to occur across species, and whether biases in morphospace correspond to phenotypes that are readily “accessible” by simple mutations within species.

Finally, our analyses shed light on biological aspects of pattern formation as well. For example, species with vertical bars tended to occupy larger regions of morphospace than species with horizontal stripes, reflecting greater variability among individuals in the former than the latter (**Fig. 2** and **Fig. 3C**). This observation suggests the possibility of a more dynamic pattern-forming process in barred species, and one perhaps less dependent on positional information in the tissue environment to set the location of initial pattern elements (e.g., interstripe iridophores of zebrafish [47]), or homeostatic interactions that maintain the integrity and position of pattern elements as fish grow (e.g., melanophore–xanthophore interactions of zebrafish [58]). Indeed, patterns in barred species appeared often to have intercalary elements, presumably reflecting transitional states.

The phenotypes we document further provide clues to the roles played by different cell types in pattern development and evolution, and the phenotypic consequences of mutations at specific loci. We confirmed and quantified roles for xanthophores in organizing melanophores across three *Danio* species. In *D. rerio, csf1ra* mutants initiate stripe formation because of interactions between interstripe iridophores and melanophores, yet xanthophores and their precursors fail to develop, so interactions required to organize and maintain stripes are missing and melanophores remain scattered [30, 39, 43, 58, 67]. In this species, we found that pattern defects were less severe anteriorly than posteriorly, perhaps reflecting an anterior–posterior progression of interstripe iridophore development, and a patterning influence of iridophores on melanophores even when xanthophores are missing [60, 68]. In *D. aesculapii* and *D*. aff. *albolineatus, csf1ra* mutants were essentially devoid of pattern, similar to *D. rerio* mutants that lack both xanthophores and iridophores (*csf1ra*; *ednrb1a*) [69]. The nearly uniform arrangement of melanophores in xanthophore-free *csf1ra* mutant *D. aesculapii* further suggests that iridophores of this species do not provide the robustness to pattern formation thought to be conferred by iridophores of *D. rerio*, based on computational analyses [35]. Whether this difference reflects a change in the strength or quality of interactions between pigment cells, or differences in the times or locations of iridophore development (*cf*. xanthophores of *D. albolineatus* [60]), remains to be determined. That regulative patterns of stripes (*D. rerio*) or bars (*D. aesculapii*) were generated in the absence of iridophores further demonstrated the sufficiency of melanophore–xanthophore interactions in driving the formation of a periodic pattern in both species. Finally, our comparison of *kita* mutant phenotypes suggested different consequences for melanophore complements, with residual melanophores occurring in both *D. rerio* and *D. aesculapii* but very few such cells in *D*. aff. *albolineatus*, despite alleles likely to have a similar abrogation of signaling through the Kita receptor tyrosine kinase. This difference in penetrance might reflect relative proliferative abilities of residual melanophores [6]. Importantly, these divergent roles for pigment cells and differences in mutational effects across species would not have been apparent without an explicitly genetic and quantitative approach, and they indicate the potential for a more complete genetic deconstruction [40] of these and other phenotypes in *Danio*.

## Materials and Methods

### Fish

All *Danio* species and mutants were maintained in standard 14L:10D at 28.5C. All analyses were conducted with approval of the University of Virginia Animal Care and Use Committee and complied with United States federal guidelines for ethical use of animals in research. *D. rerio* were wild-type AB^wp^, *csf1ra*^*j4e1*^ [42] *kita*^*b5*^ [45], and *ltk*^*j9s1*^ [46]. Other *Danio* species were obtained from tropical fish suppliers and have been maintained in the lab under conditions similar to *D. rerio*. Mutants of *D. aesculapii* and *D*. aff. *albolineatus* were generated by CRISPR/ Cas9 mutagenesis and isolated as stable lines by standard methods [70, 71]. To target *csf1r* in *D. aesculapii* and *D*. aff. *albolineatus* a single target site in exon 2 was selected (GGATCAGGACACCCTTTCTG). Two *kita* target sites in exon 2 were used for *D. aesculapii* (GGGAAAATATTCATGCCGAG; GGACCTTGTGGGGTAATGGT) and two *kita* target sites in exon 3 were selected for *D*. aff. *albolineatus* (GGTTCAAGTCTTTCATATCT; GGCGGTGGAAAAAGTCAGGA). To generate a viable allele of *ltk* in *D. aesculapii* we targeted a site corresponding to *D. rerio ltk*^*j9s1*^ by homology directed repair [72, 73]. An oligonucleotide (AGCAGATGGACAAGATGGCCTCTCTTTTGTTCACCCCATGGGAAAGATATTCCTCCAGTCTTTAGCTGGTCAGACTTAACCCAATCTTGACTATGTATAGTGATGTTGACTTGTACTTGT) encoding the same missense allele as present in *ltk*^*j9s1*^ of *D. rerio* was coinjected with CRISPR/ Cas9 reagents; a majority of resulting *D. aesculapii* alleles exhibited the intended lesion and all individuals included in analyses lacked iridophores. Alleles are shown in **Fig. S5**.

### Imaging

Adults were imaged using Nikon D200 or D810 single lens reflex cameras with 105 mm f2.8 IF-ED Micro-Nikkor lens. Approximately equal numbers of males and females of eleven species were included: *D. rerio, n*=11, *D. rerio csf1ra, n*=9, *D. rerio kita, n*=10, *D. rerio ltk, n*=10; *D. aesculapii, n*=15, *D. aesculapii csf1ra, n*=12, *D. aesculapii kita, n*=10, *D. aesculapii ltk, n*=9; *D. quagga, n*=15; *D. kyathit, n*=9; *D. nigrofasciatus, n*=14, *D. tinwini, n*=12; *D*. aff. *albolineatus, n*=14, *D*. aff. *albolineatus csf1ra, n*=9, *D*. aff. *albolineatus kita, n*=9; *D. kerri, n*=9; *D. choprae, n*=19, *D. erythromicron, n*=11; *D. margaritatus, n*=8.

### Pattern analyses

Initial processing of images for pattern analyses were performed in Adobe Photoshop CC. RGB color images were converted to greyscale and pixel intensities adjusted linearly so that background whites (plastic ruler in each image) were set to 255 and blacks (within pupil of eye) were set to 0. Regions of interest (ROIs) were defined with proportions of 2 x 1 (length x height). ROI height and dorsoventral position were set to correspond with lines drawn anteriorly from the tail at the caudal peduncle (base of caudal fin). ROI width was then set to twice this height, and the posterior edge of the ROI was defined to correspond with a point just anterior to the posterior margin of the dorsal fin insertion. Images were then cropped to 1000 x 500 pixels for analysis. Entropy and FFT analyses did not differ markedly when using either of two alternative ROIs (anatomically defined as above, but with proportions of 3 x 1; or having fixed dimensions rather than scaled to size, with proportions of 2 x 1).

For entropy analyses of each image, the entropy (H) for each row (*Z*_row_) or column (*Z*_col_) was calculated [48]:

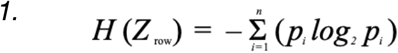

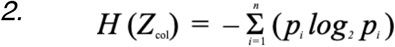

where *n* is the number of possible states of *Z* and *p*_*i*_ is the probability of finding the i^th^ state of *Z*. For each image, mean entropies were calculated for: (1) all rows (*E*_*DV*_); and (2) all columns (*E*_*AP*_). Entropy analyses were conducted using MATLAB R2021b (MathWorks, Natick, MA, USA). Preliminary analyses indicated that application of a Gaussian blur (20 pixels in Adobe Photoshop CC) removed variation from fish to fish associated with degree of pigment dispersion within cells while preserving overall pattern attributes; final entropy analyses employed these adjusted images.

For FFT analyses, grayscale ROI images were imported into MATLAB and rescaled to 500 × 500 pixels. The two-dimensional discrete Fourier transformation of the pixel matrix was calculated using MATLAB’s native 2-D FFT algorithm. The magnitude component of the transformation contains the geometric patterning information of an image, and brightfield images are predominantly represented by low-frequency data [51, 74, 75]. The magnitude component was retained and the matrix was shifted so that zero-frequency components were centered. To obtain values of pattern orientation (*Ω*) and strength (*ψ*), FFT output matrices (*a*) were cropped to 21 × 21 pixels and contained the lowest frequency data for analysis. As pattern changes vertically and horizontally were recorded across the X and Y axes of the centered matrix, the ratio of the sum of the squares Y to X values across the five centermost rows or columns (equations *3*–*5*) were used to assess pattern orientation [51, 75]:

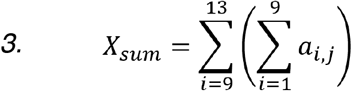

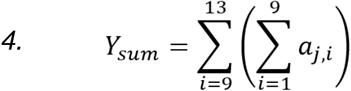

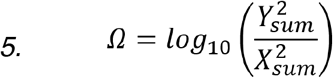

Magnitude values at individual frequencies reflect the extent of pattern change at that frequency [51, 75], therefore the strength of the pattern can be evaluated independent of the orientation of the pattern. The strength of pattern score *ψ* was calculated as the natural logarithm of the total sum of the same X and Y values generated for the orientation calculation, normalized by the number of pixels and their intensity range, then multiplied by 100 to fit within an order of magnitude the scale of *Ω* (equation *6*):

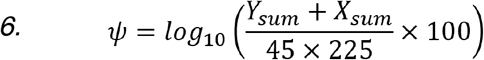

In addition to entropy and FFT analyses, pattern variation was measured as previously described [38]. The blurred grayscale images used for entropy analyses were thresholded to segment the patterns into melanophore elements and background by comparison to brightfield images. Arrays containing the proportion of pixels belonging to a melanophore pattern element at each position along the AP and DV axes was determined using the average 8-bit gray values along each axis. Stripe variation (*E*_*DV*_) and bar variation (*E*_*AP*_) were calculated as the standard deviation of these arrays.

### Ancestral state reconstruction

Ancestral state reconstruction of both discrete and continuous characters was performed on a previously published phylogeny based on reduced-representation sequencing [14]. Discrete character reconstructions were estimated using the ace() function from the ape package with an All Rates Different model specified [76].

### Statistics

Analyses were performed using JMP Pro 16 (SAS Institute, Cary NC) for Macintosh. In analyses of variance and *t*-tests, residuals were inspected and variables log-transformed or square root-transformed to restore normality or homoscedasticity. Discriminant analyses used quadratic fitting to allow for differences in within-group covariance matrices.

## Abbreviations

Csf1ra: Colony stimulating factor 1 receptor a;
*E*_*AP*_: bar entropy;
*E*_*DV*_: stripe entropy;
FFT: Fast Fourier Transform;
Kita: Kit receptor tyrosine kinase a;
Ltk: Leucocyte tyrosine kinase;
ROI: region of interest;
*V*_*AP*_: bar variation;
*V*_*DV*_: stripe variation;
*Ω*: pattern orientation;
*ψ*: pattern strength;

## Acknowledgements

Thanks to Amber Schwindling and other Parichy lab members for helping to maintain the fish and to Mike Nishizaki for assistance with FFT analyses.

## Author contributions

Conceptualization: DMP, LBP; mutants and imaging: LBP, YL, VML; development of quantitative metrics, image analysis, statistics: BMM, DMP, VML; drafting of manuscript: DMP, LBP, BMM.

---

**Fig. S1.**
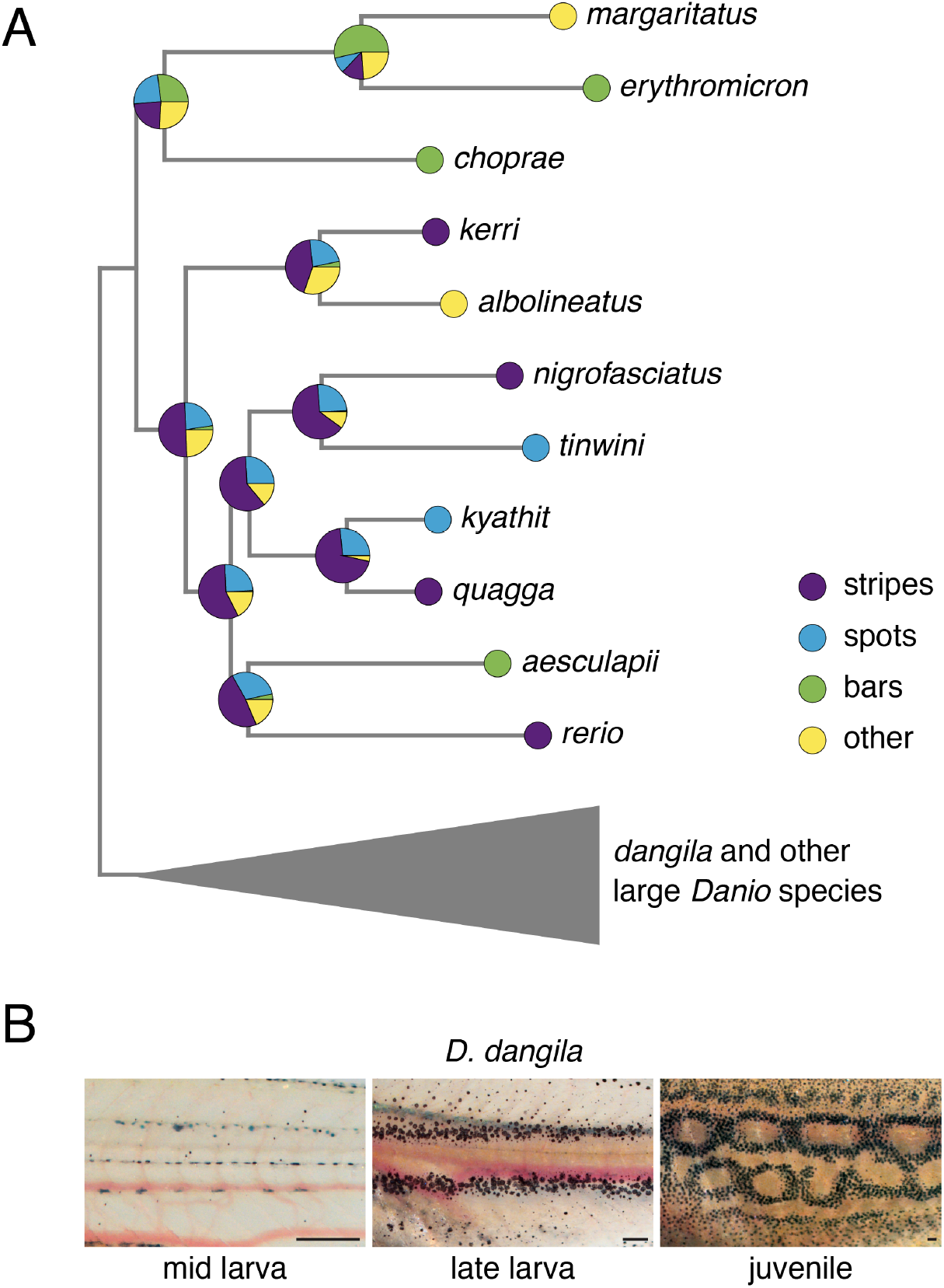
Ancestral state reconstruction of *Danio* patterns as a discrete character. (A) Phylogeny of small *Danio* species with ancestral state probabilities under the maximum likelihood model allowing for different rates of pattern state transitions. The position of large *Danio* species as outgroup is shown, but the uncertain relationships within this group preclude it from properly informing the ancestral state at the base of small *Danio* species. (B) Pattern development in *D. dangila*. The chain motif shared by many large *Danio* species emerges from a striped juvenile pattern very similar to *D. rerio*.

**Fig. S2.**
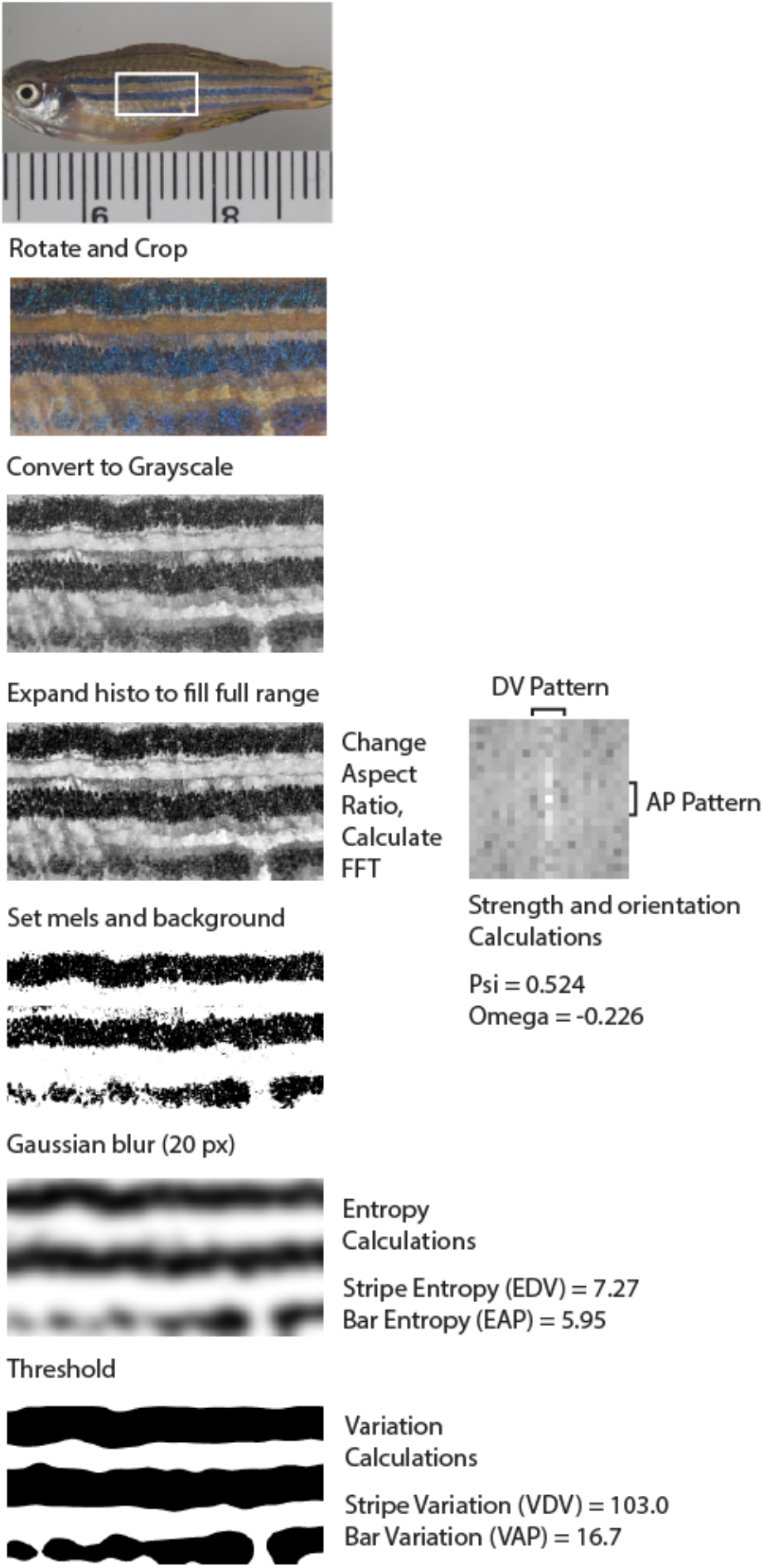
Image processing and analysis pipeline. Brightfield images are cropped to a 2:1 region of interest near the center of the fish. RGB images are corrected for background and converted to grayscale. Histograms are stretched to account for uneven lighting between fish and these images are used for FFT analyses. High contrast images are created by setting the lightest melanophores to black and the darkest background to white. High contrast images are blurred and these images are used for entropy calculations. Blurred images are compared to brightfield images and thresholded to give segmented, binary patterns for variation calculations.

**Fig. S3.**
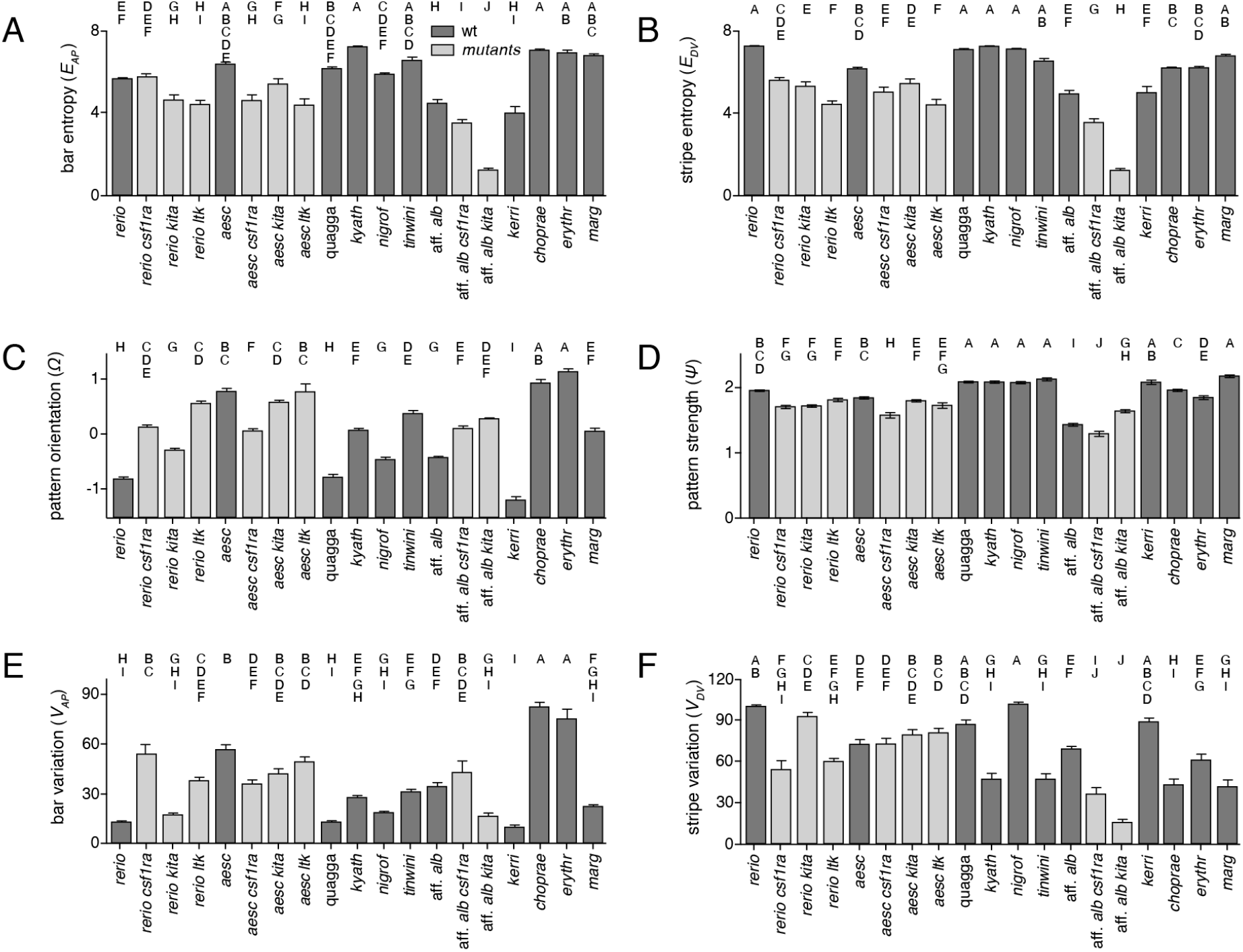
Pattern metrics used in analyses. Shown are univariate plots (means±SE) for pattern metrics described in the main text. Dark grey bars indicate wild-types, light grey bars indicate *csf1ra, kita*, and *ltk* mutants. Shared letters above bars indicate means that were not significantly different (*P*>0.05) in Tukey-Kramer post hoc comparisons (all ANOVA, *P*<0.0001).

**Fig. S4.**
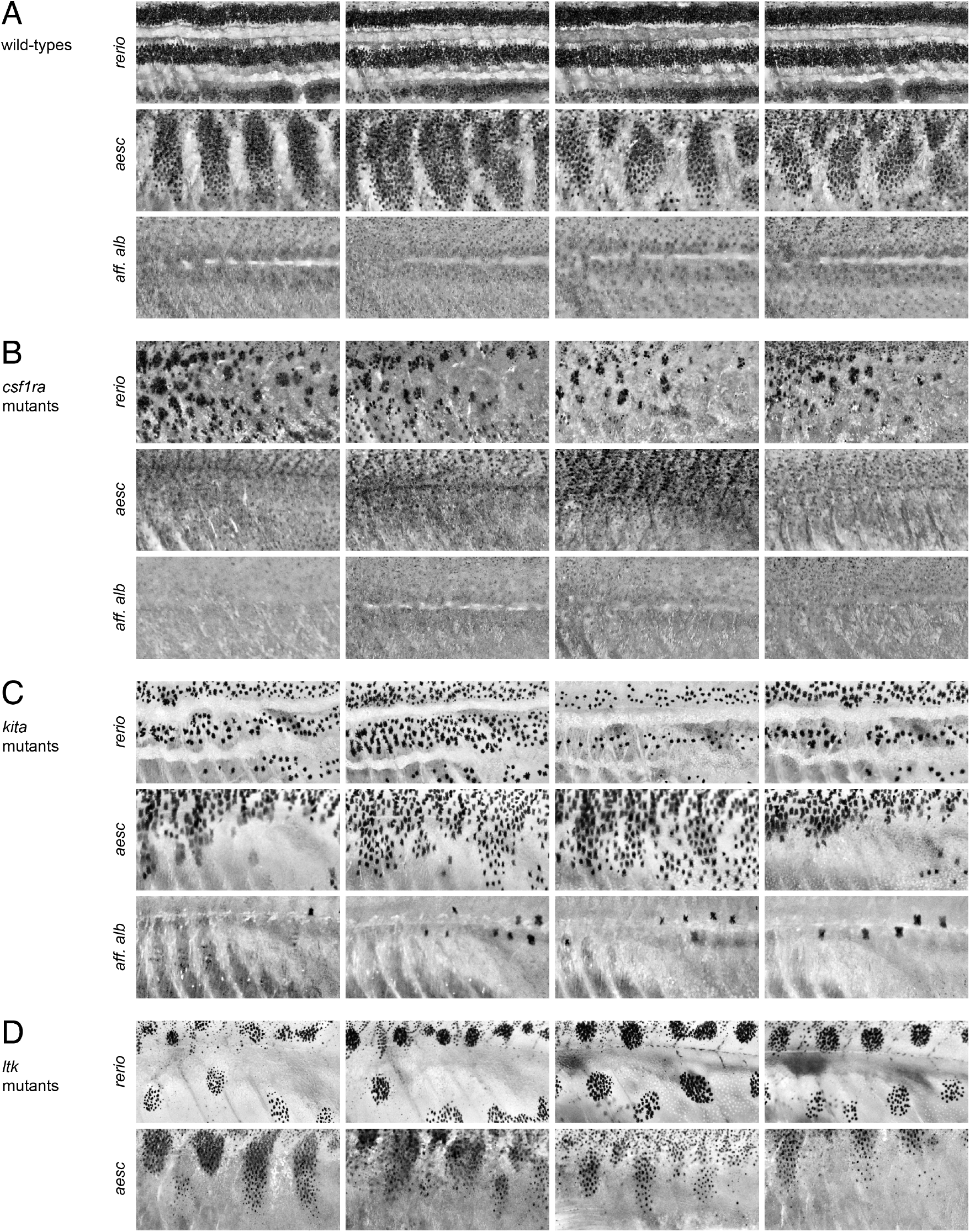
Mutant phenotypes of *Danio* species. Representative individuals are shown, illustrating typical variation among individuals. Left two panels, males; right two panels, females. (A) Wild-types of each species (same individuals shown in Fig. 2, provided here for ease of comparison). (B) *csf1ra* mutants lacking xanthophores. (C) *kita* mutants deficient for melanophores. (D) *ltk* mutants lacking iridophores.

**Fig. S5.**
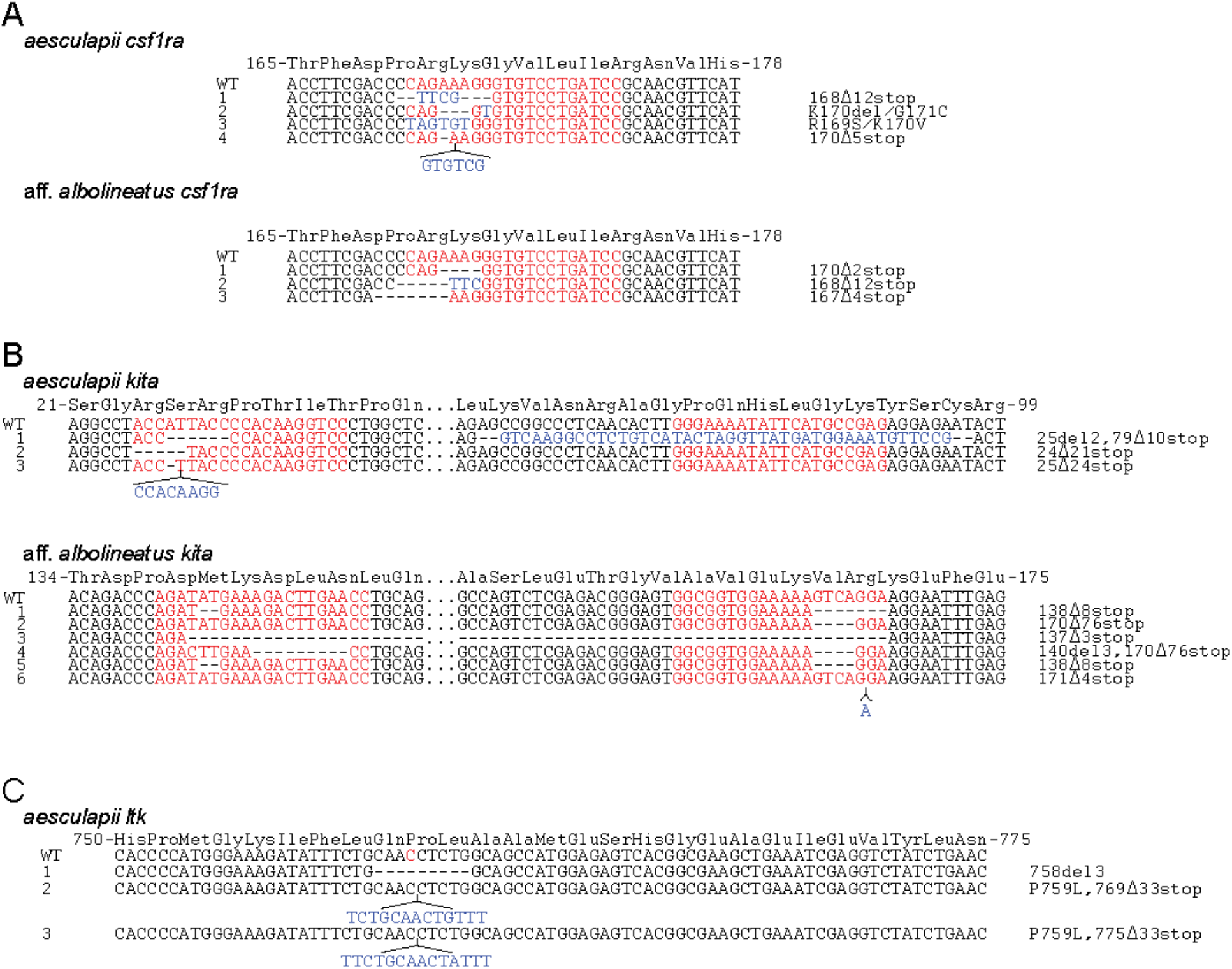
Sequences of mutant alleles identified in fish used for pattern analyses. (A) *csf1ra* mutant alleles. (B) *kita* mutant alleles. (C) *ltk* mutant alleles. Red letters indicate CRISPR target sites in (A) and (B), and the site corresponding to *D. rerio ltk*^*j9s1*^ in (C). Deletions (–) and insertions or small duplications (blue) are indicated in mutants. Designations indicate first affected amino acid and number of deleted (del) or altered (Δ) amino acids prior to premature stop codon if present, or changes in specific amino acids.

**Table S1.**
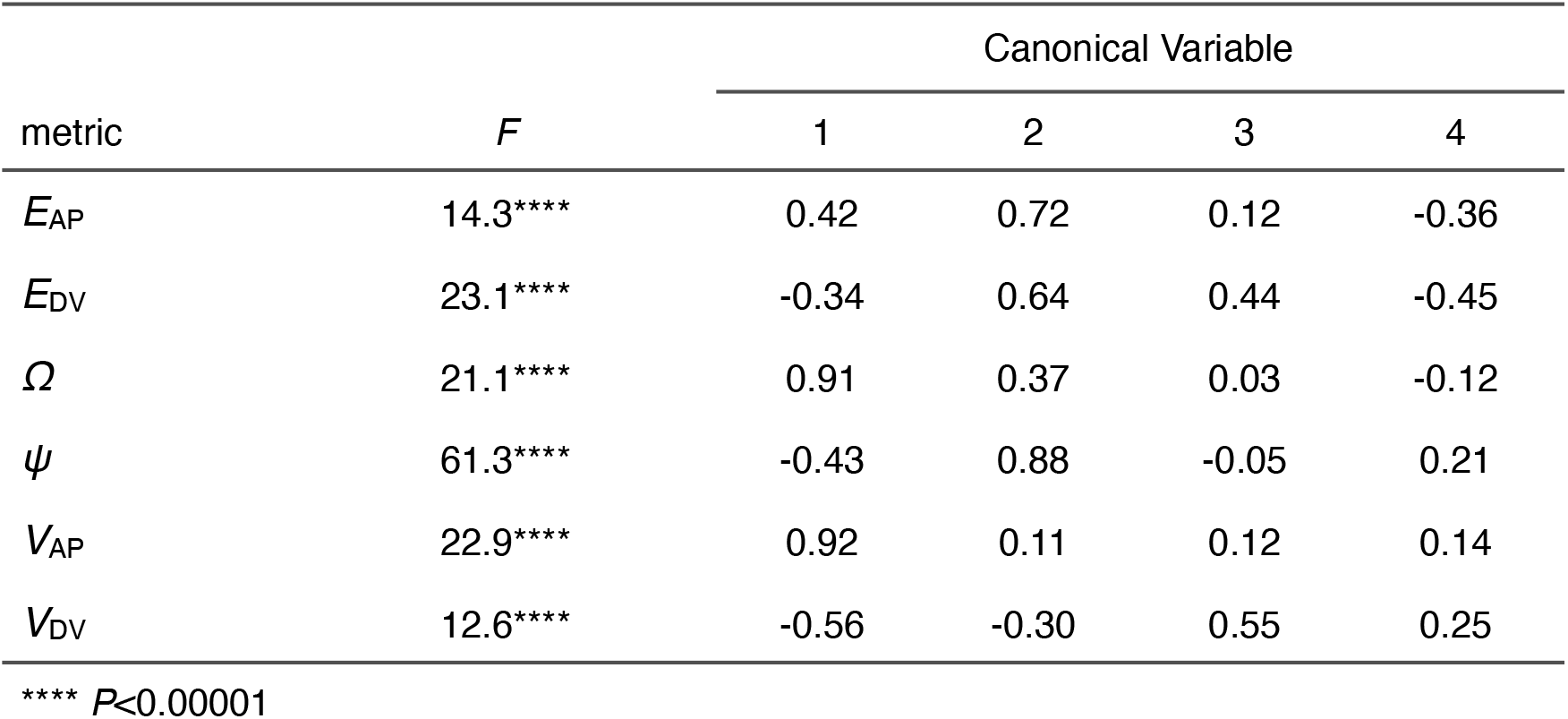
Multidimensional pattern space. Output from discriminant analysis of wild-type individuals using six pattern metrics and quadratic fitting, indicating individual factor significance when tested against full model (*F* statistic) and standardized loadings for canonical variables 1–4.

**Table S2.** Bivariate correlations among pattern metrics. *F*_1,165_ statistics are shown above diagonal and *R*^2^ values are given below diagonal. Fitted models included wild-types of all species and all available *csf1ra, kita*, and *ltk* mutants.

| metric | $E_{AP}$ | $E_{DV}$ | $\Omega$ | $\psi$ | $V_{AP}$ | $V_{DV}$ |
| --- | --- | --- | --- | --- | --- | --- |
| $E_{AP}$ | — | 834.4*** | 16.1*** | 104.8*** | 35.7*** | 0.1 |
| $E_{DV}$ | 0.80 | — | 3.8 | 152.2*** | 0.0 | 33.1*** |
| $\Omega$ | 0.07 | 0.02 | — | 2.6 | 325.1*** | 71.6*** |
| $\psi$ | 0.33 | 0.42 | 0.01 | — | 3.8 | 4.6* |
| $V_{AP}$ | 0.14 | 0.00 | 0.60 | 0.02 | — | 42.7*** |
| $V_{DV}$ | 0.00 | 0.13 | 0.25 | 0.02 | 0.17 | — |

\*\*\* $P < 0.0001$ , \*\* $P < 0.001$ , \* $P < 0.05$

**Table S3.** Effects of sex differences on pattern metrics. Shown are results of ANOVAs for each metric that considered effects of species and genotype (combined for simplicity), sex, and interactions between sex and species–genotype. Right columns indicate variance explained (adjusted *R*^2^) by models that included sex and specifies–genotype interaction with sex (“full”) and models without these factors (“species–genotype only”). Effects of sex varied among species (“sex x species–genotype”) but differences were minor and accounted for ≤4% of overall variances.

| metric | species–<br>genotype<br>$F_{37,177}$ | sex<br>$F_{1,177}$ | sex x species–<br>genotype<br>$F_{37,177}$ | $R^2$<br>(full) | $R^2$<br>(species–<br>genotype only) |
| --- | --- | --- | --- | --- | --- |
| $E_{AP}$ | 80.2*** | 10.1** | 4.0*** | 0.90 | 0.86 |
| $E_{DV}$ | 96.5*** | 5.6* | 2.8** | 0.92 | 0.89 |
| $\Omega$ | 140.9*** | 3.5 | 3.4*** | 0.94 | 0.92 |
| $\psi$ | 105.3*** | 9.5* | 3.0*** | 0.93 | 0.90 |
| $V_{AP}$ | 58.7*** | 1.4 | 3.3*** | 0.87 | 0.83 |
| $V_{DV}$ | 38.2*** | 4.1* | 1.8* | 0.81 | 0.77 |

\*\*\* $P < 0.0001$ , \*\* $P < 0.001$ , \* $P < 0.05$

